# EMinsight: a tool to capture cryoEM microscope configuration and experimental outcomes for analysis and deposition

**DOI:** 10.1101/2024.02.12.579963

**Authors:** Daniel Hatton, Jaehoon Cha, Stephen Riggs, Peter J. Harrison, Jeyan Thiyagalingam, Daniel K. Clare, Kyle L. Morris

## Abstract

The widespread adoption of cryoEM technologies for structural biology has pushed the discipline to new frontiers. A significant worldwide effort has refined the Single Particle Analysis (SPA) workflow into a reasonably standardised procedure. Significant investment of development time have been made particularly in sample preparation, microscope data collection efficiency, pipeline analyses and data archiving. The widespread adoption of specific commercial microscopes, software for controlling them and best practises developed at national facilities has also begun to establish a degree of standardisation to data structures coming from the SPA workflow. There is opportunity to capitalise on this moment in the field’s maturation, to capture metadata from SPA experiments and correlate this with experimental outcomes, which is presented here in a set of programmes called EMinsight. This tool aims to prototype the framework and types of analyses that could lead to new insights into optimal microscope configurations as well as for defining methods for metadata capture to assist with archiving of cryoEM SPA data. We also envisage this tool to be useful to microscope operators and facilities looking to rapidly generate reports on SPA data collection and screening sessions.

**Synopsis:** EMinsight is a Python-based tool for systematically mining metadata from single particle analysis cryoEM experiments. The capture and analysis of metadata facilitates assessment of instrument performance, provides concise reporting of experiment performance and sample quality by analysing preprocessing results, and gathers metadata for deposition. We envisage this approach to benefit the microscope operator, facility managers, database developers and users.

## 1. Introduction

Cryogenic-sample electron microscopy (CryoEM) has undergone significant growth and has matured into a major tool for determining the structures of macromolecular complexes at resolutions useful in structural biology research. This progress is evident in the substantial number of entries in the Electron Microscopy Data Bank (EMDB) which at the time of writing stands at 24,576 for single-particle analysis (SPA). The SPA technique concerns using a transmission electron microscope (TEM) instrument to acquire many thousands of 2-dimensional images of a target biological macromolecule preserved under cryogenic conditions and using computational techniques to identify the different poses of that macromolecule to reconstruct its 3-dimensional structure. The growth of the technique can be attributed to improvements in various aspects of the SPA workflow, including sample preparation, automation and efficiency gains in data collection, and data analysis techniques.

One crucial aspect that has gained prominence in enhancing microscope data collection efficiency is the computerised control of microscopes. Several software packages, such as Leginon (Carragher *et al*., 2000), SerialEM (Mastronarde, 2003), and Thermo Fisher Scientific’s (TFS) EPU (E Pluribus Unum – Out of Many, One), have emerged as major tools in instrument control allowing increased levels of data production via autonomy and lowering the technical barrier in controlling TEMs. At the time of writing, 72% (17,657) of the SPA macromolecular structures deposited in the EMDB are recorded as having been performed on a Titan Krios microscope, as now produced by Thermo Fisher Scientific, reflective of a standardisation that has occurred due to the predominance of an instrument type in the field. The evident widespread adoption of instrumentation and collection strategies presents an opportunity to develop processes that attempt to standardise the capture of metadata from cryoEM imaging experiments. Such a process would benefit the community, enabling the automatic and robust generation of descriptions of how an experiment was performed. Such a tool to survey or parse instrument metadata also presents the opportunity for facilities to globally monitor the utilisation and performance of their instruments.

Figure 1 graphically represents the SPA workflow at (a) the level of the experiment and (b) in image processing. Acquisition workflows are reasonably well standardised in SPA. Since the images (or micrographs) taken of the target macromolecules are destructive due to radiation damage, these data may only be collected from an area once. Thus, the experimental workflow may be thought of as a targeting exercise, whereby an expert operator uses non-destructive low-dose low-magnification images to identify regions of the specimen (Atlas and Grid Square) expected to yield data of high quality. The knowledge of what regions produce high-quality data may have been established from prior knowledge gained during trial collections on the current or equivalent specimen (so called “screening”) but in essence the operators goal is to set the microscope to target the co-ordinates of many Foil Holes and within those, many Acquisition Areas. The microscope will then automatically collect Micrograph Data in those Acquisition Areas. This targeting exercise collects and produces a hierarchical image structure where acquired micrographs exist in relation to a series a lower magnification images that describe that micrograph’s location on the specimen.

**Figure 1.**
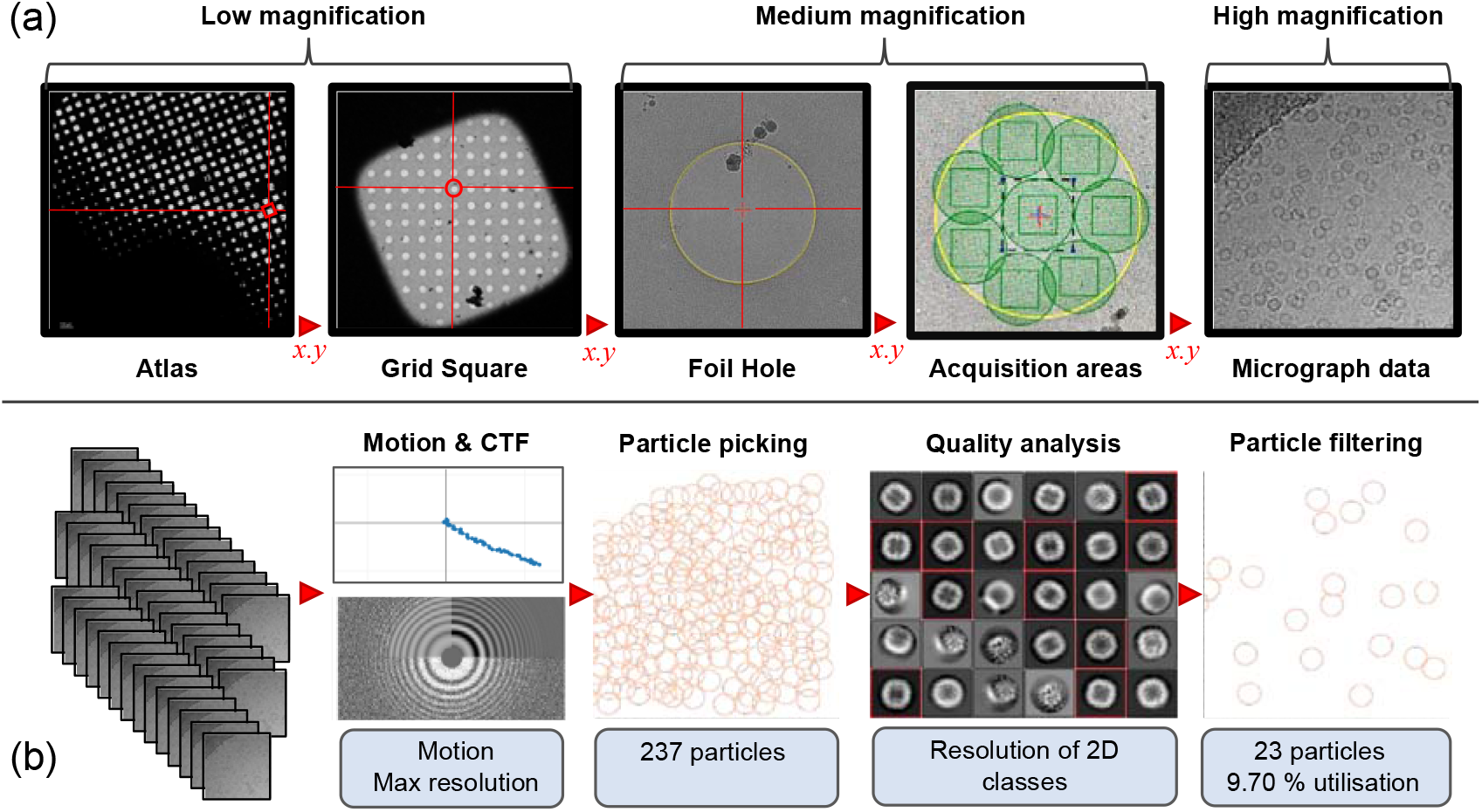
Workflow for an SPA cryoEM experiment showing (a) the hierarchical image collection structure representing the collection strategy to arrive at acquiring micrograph data and (b) the preprocessing of many micrographs to arrive at filtered particles ready for structure determination. Some potential quality metrics that are obtained from preprocessing are highlighted in light blue.

Assessments of the quality of micrographs from SPA experiments are made as a product of the image processing that is performed to transform 2-dimensional images into a 3-dimensional structure of the target macromolecule. This is described in detail in various general reviews (Orlova & Saibil, 2011, Saibil, 2022, Lyumkis, 2019). Due to the broad range of softwares available for structure determination in SPA cryoEM, image processing workflows can uniquely evolve for each structure determination project. Increasingly however, in-line analysis packages are available to perform the image processing steps leading up to particle identification (so called particle picking) and 2D alignment, averaging and classification (Punjani *et al*., 2017, Fernandez-Leiro & Scheres, 2017, Gómez-Blanco *et al*., 2018, Tegunov & Cramer, 2019, Caesar *et al*., 2020) which we refer to as a preprocessing pipeline and depicted in Figure 1b. Many packages able to automatically perform analyses to produce 3-dimensional reconstructions and the quality of the 3-dimensional density is the defacto measure of the experiments success, however preprocessing pipelines arguably already produce many of the metrics suitable for describing micrograph data quality. Benefitting again from a relative standardisation of the preprocessing approach there is an opportunity to capture these quality metrics as metadata describing the quality of an acquired dataset of micrographs.

Taken together these metadata describe how the experiment was configured and performed, and the experimental and analytical outcomes relating to the instrument and specimen performance. Many of these metadata points may still be manually documented by users but could equally be retrieved from outputs from the instrumentation and pipelines that performed and analysed the experiment. If performed automatically this would lead to efficiency gains for depositors and an increase in the robustness of the deposition process. Automatic capture of metadata describing the cryoEM experiment could also lower the barrier to capturing and depositing more descriptive data on the experiment. If available this would permit hypothesis testing on the relationship between experimental configuration and experimental outcome. We would expect an increased richness of metadata in the structural biology EM archives (Lawson *et al*., 2016, Iudin *et al*., 2023, Patwardhan & Lawson, 2016) will also make entries ready to support future machine learning (ML) applications that require more descriptive labels for training and inference.

We present a tool, called EMinsight, which allows the systematic extraction of information from TFS EPU SPA directories to collate and summarise metadata describing the experiment. The directories of associated pipeline preprocessing in Relion are also interrogated to associate quality information with a collected dataset. The outputs of EMinsight are expected to be useful to different ends users: PDF reports on the experiment for the microscope operators documentation, comprehensive metadata capture for facility database and instrument managers, and concise metadata capture with data integrity measures for archive and deposition developers. We expect this type of tool can form the conceptual basis for future systems that could be used locally within facilities to monitor instrument and session performance. Additionally, EMinsight could represent one type of approach to a method of generating metadata to automatically populate archival depositions of macromolecular structures, maps and raw data of SPA cryoEM experiments.

## 2. Materials and methods

### 2.1. Data structure

EMinsight has been developed with the SPA session type performed at the UK national cryoEM facility eBIC, Diamond. The software expects a data structure as shown in Table 1.

**Table 1.**
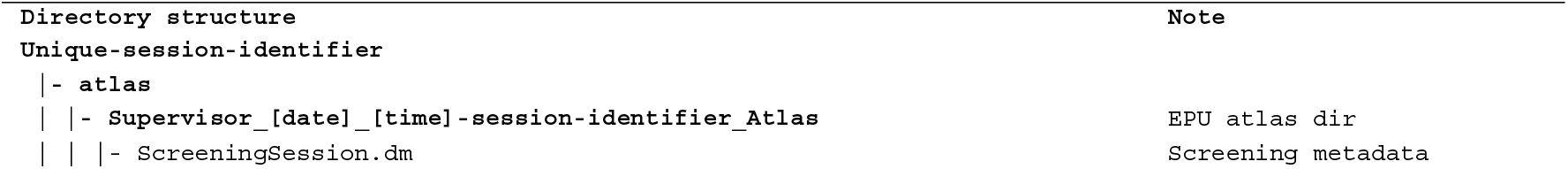
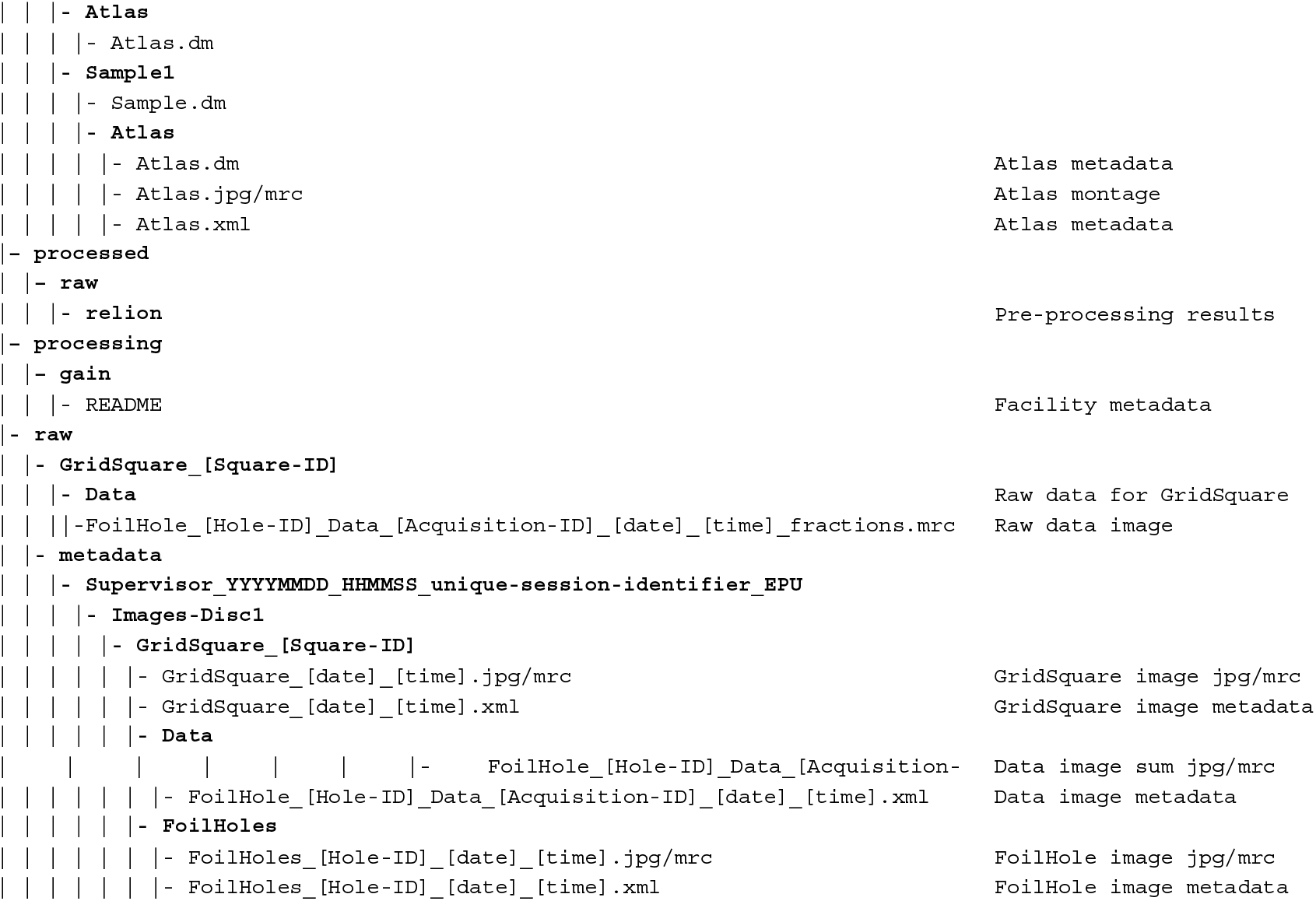
A representation of the directory data structure expected by EMinsight.

### 2.2. XML metadata parsing

Under the SPA collection system EPU (Thermo Fisher Scientific) metadata describing the experiment is stored in XML format. These files are produced by EPU during an SPA session and store much of the metadata describing the microscopes configuration and behaviour during the experiment. EMinsight has been developed at eBIC and generally across EPU versions 2 and 3, suggesting a degree of stability in the format of these files produced by the instrument manufacturers software. In general, every image that is acquired by the microscope and collection system is paired with an XML metadata file documenting the optical configuration of the microscope for that image as well as additional information. Information about the setup of the experiment is stored in hierarchical auxiliary files that do not necessarily have images associated with them (also in XML format file, with the extension DM). For instance, the number and names of grids inventoried for collection is stored in a global ScreeningSession.dm but the individual grid type, hole diameter and radius are stored in each EpuSession.dm for each grid. EMinsight uses a python library to transform the XML into a dictionary and then query known addresses for instrument and experimental metadata. In general, these are passed to the pandas python library to create internal dataframes for reference, analysis and output.

### 2.3. Preprocessing analysis parsing

During SPA collections at eBIC, an automatic preprocessing pipeline is executed using RELION and the CCP-EM pipeliner. This includes motion correction, CTF estimation, particle picking (crYOLO), 2D classification, particle selection and subsetting. The directory structure of results is as expected for RELION v4 but also includes a relion_it_options.py file containing many of the parameters defining the processing pipeline. EMinsight parses the pipeline parameters and results to gather these data and associate them with each individual micrograph of a dataset. Associations of those micrographs to their respective originating grid square and hole locations are retained such that they can be used for data grouping in location-based analyses.

### 2.4. Particle picking analyses

Particle picking is typically performed at eBIC using crYOLO (Wagner *et al*., 2019). This is leveraged to read the particle diameter by parsing and averaging the particle diameters reported for each picked particle by crYOLO in the output ^*^.cbox files. Particle density is calculated as a value normalised to 1, where 1 would be maximum packing of packing into a simplified 2D array in the micrograph field of view based on the observed particle diameter. Particle coordinates are clustered using the SciKit-learn implementation of nearest neighbour analysis. Where the nearest neighbour distance is found to be less than 80% of the measured particle diameter, that particle is labelled as overlapping or as termed in EMinsight, clustered.

### 2.5. Outputs

Each of the following subsections describes the outputs the a user of EMinsight can expect to be produced.

#### 2.5.1. Comma separated value (csv) collated data

^*^_datastructure.csv: associates the micrographs with associated lower magnification images, as well as quality metrics gathered from metadata.

^*^_optics.csv: reports all of the microscope optics configurations for each preset used by EPU for the data collection session.

^*^_processed.csv: reports on the outcomes of pre-processing jobs to infer quality of the dataset.

^*^_session.csv: reports on the outcomes of the session, i.e. targeting statistics, collection rates, dataset size, specimen properties, collection strategy.

#### 2.5.2. Reports

PDF reports are created to summarise and present the major descriptors of the data collection session to the end user of EMinsight. These are ^*^_session.pdf to present information on the session configuration and outcome and ^*^_processed.pdf to present information on the pre-processing outcomes.

#### 2.5.3 Deposition

JSON files recording the necessary fields for populating the Microscopy Section of an EMDB deposition. Checksums are included to provide a method of verifying data integrity.

## 3. Results

### 3.1. Reporting on SPA sessions

All raw data and the hierarchical image structure from SPA experiments are written out by TFS EPU along with metadata in XML format. These can be interrogated to expose the configuration of the experiment, however these metadata are not practically human readable. EMinsight parses through the metadata structure of experimental outputs from Thermo Fisher Scientific EPU sessions to gather important data describing the experiment to produce a concise human readable report. EMinsight can be executed from the command line or a simple user interface, as shown in Figure 2.

**Figure 2.**
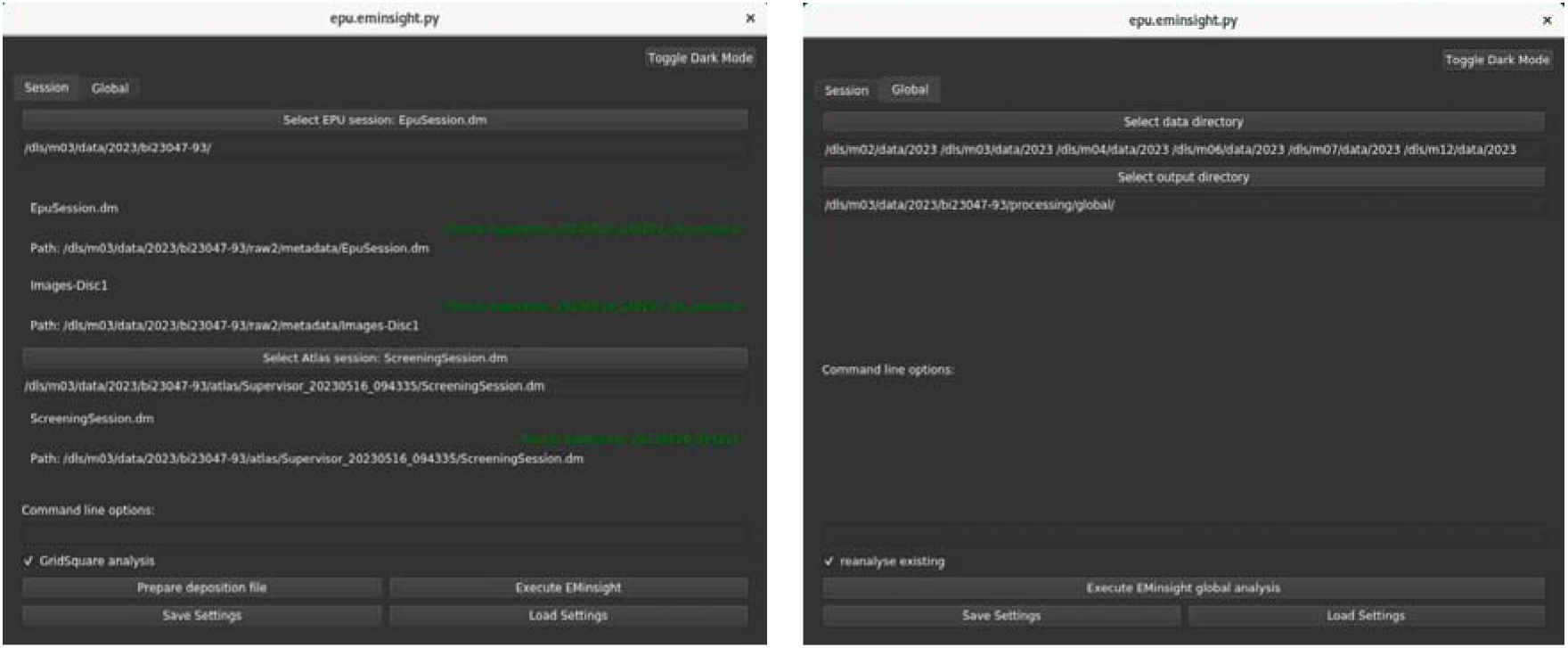
The EMinsight user interface, (left) for analysing a single cryoEM SPA data collection session and (right) for analysing and collating session analyses across multiple directories from a SPA user programme.

Example PDF reports are shown in Supplemental Information S1 and are expected to be useful as reference documents when stored as part of an electronic lab notebook by the microscope operator and EMinsight user. The types of metadata that are captured and exposed by EMinsight are summarised in Table 2. In addition to capturing Instrument Configuration metadata, additional properties of an SPA session may be described as Experimental Outcomes, Calculated Parameters and Analytical Outcomes. Many of these descriptors are displayed in the PDF reports but all exposed descriptors are written to csv files serving as a simple collated data for each session as found in Supplemental Information S2.1. EMinsight is further capable of parsing multiple experiments and globally aggregating collated data as found in Supplemental Information S2.2.

**Table 2.** A selection of the metadata captured and exposed by EMinsight to the user.

| Instrument Configuration | Experimental Outcomes | Calculated Parameters | Analytical Outcomes |
| --- | --- | --- | --- |
| C2 | Total available squares | Dose rate ($e^-/\text{\AA}^2/\text{s}$) | Motion (early/late/total) |
| Spot size | Targeted squares | Rate (mic/hr) | CTF resolution (min/max) |
| Beam size ($\mu\text{m}$) | Collected squares | Rate ($\mu\text{m/hr}$) | Particle size |
| Magnification (X) | Collected images |  | Particle count |
| Defocus range ($\mu\text{m}$) | Collected area ($\mu\text{m}$) | | Particle density |
| Grid type | Acquisition time stamps |  | Particle clustering |
| Hole size/spacing ($\mu\text{m}$) | Acquisition locations | | Class assignment resolution |
| Shots per hole | Acquisition data structure |  |  |

### 3.2. Instrument performance assessments

EMinsight performs systematic analyses on individual data collection sessions as well as comparing these analyses across multiple sessions from a cryoEM instrument as part of a facility or user programme. In one example, this type of analysis confirms the speed gains that can be attained on an eBIC microscope (TFS Titan Krios) by increasing magnification and employing multi shot collection facilitated by aberration free image shifting (AFIS) multi shot. However, it is important to note that these speed gains are not sufficient to offset the loss of field of view incurred due to the magnification increase (Figure 3a). A systematic analysis of collection rates across multiple Titan Krios instruments confirms this behaviour (Figure 3b) where multiple strategies may have been employed to enhance collection rates but are still unable to collect the same amount of usable area as magnification is increased. In light of this, microscope operators might first consider what resolution they need to achieve, using the largest pixel size appropriate for this and then carefully assessing how long they need to collect given expected data collection rates at a particular magnification and experimental set up. Considering efficient data collection using larger pixel sizes has been suggested elsewhere (Harrison *et al*., 2023). EMinsight provides a means to empirically assess what data collection rates can be expected on an instrument in a particular configuration allowing these calculations to be driven by historical performance data. Indeed a Titan Krios that has a significant configuration difference in its detection system performs differently (see Supplemental Information Figure S1) to the analyses presented in Figure 3. As vendors improve instrument and data collection workflows further speed gains may be realised. In the meantime the community may want to consider utilising the speed and area gains achievable from low magnification collection (i.e. 1.5 Å/px, superresolution 0.75 Å/px) in combination with super resolution camera detection modes to allow recapitulating high-resolution information.

**Figure 3.**
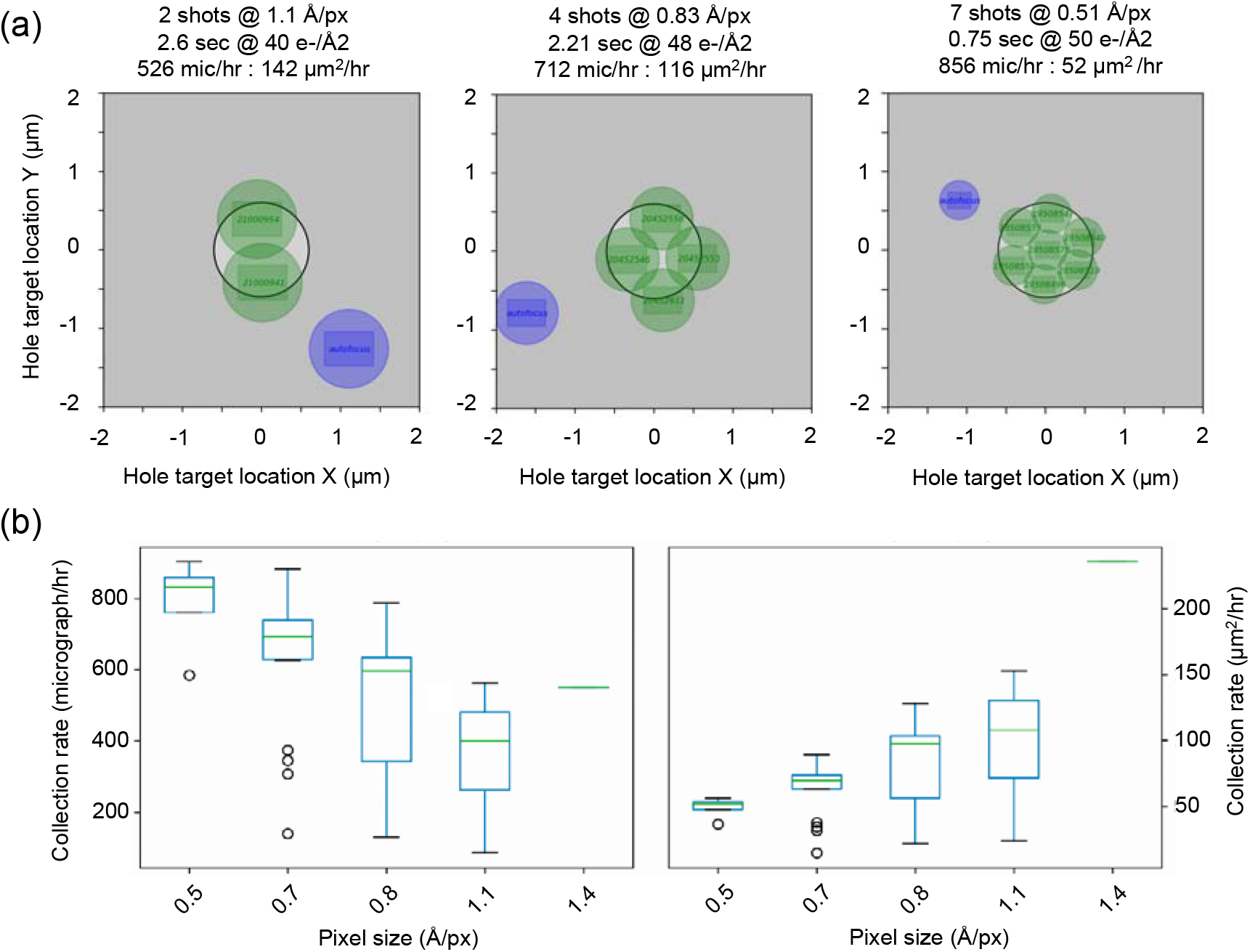
Multishot data acquisition approaches achievable by increasing microscope magnification. Representations are shown (a) with their sessions associated collection performance on a Titan Krios equipped with Gatan K3 + Bio Quantum camera/filter system. (b) The collection performance in mic/hr and um^2^/hr for the same Titan Krios microscopes with Gatan K3/bioquantum camera/filter systems is expressed in boxplots robustly revealing the speed gains obtained from increasing magnification do not offset the loss in field of view.

### 3.3 Experimental performance assessments

Where automatic processing pipelines are increasingly adopted at cryoEM facilities, it is possible to rapidly inform the operator of the quality of their sample (Punjani *et al*., 2017, Fernandez-Leiro & Scheres, 2017, Gómez-Blanco *et al*., 2018, Caesar *et al*., 2020), and critically to provide feedback with increasing detail and as early as possible during experimental time. Pipeline implementations may be customised by the facility to suit local compute infrastructure but off the shelf solutions are publicly available in packages such as RELION (Fernandez-Leiro & Scheres, 2017), CryoSPARC (Live) (Punjani *et al*., 2017) and WARP (Tegunov & Cramer, 2019). Many of these pipelines attempt to perform analyses all the way to a 3-dimensional reconstruction without user intervention, however pre-processing pipelines are more commonplace in facilities. In this manuscript, reference is made to a pipeline that performs pre-processing steps including motion correction, CTF estimation, particle picking and initial 2D classification and averaging. Figure 4 shows typical picked particle coordinates for a micrograph, as shown to the user in the report and frequently reported by common image processing softwares. EMinsight additionally performs a density and clustering analysis, as well as labelling each particle with the resolution of the 2D class it is assigned to during preprocessing. We note that the crYOLO particle picker used already excludes some particle picks found in clusters and so cases of severe particle clustering may be underestimated. These particle quality metrics can be averaged to describe a micrograph quality in a singular value. Whilst this averaging may hide subtle trends in the data, it is expected that these metrics will be useful for reporting global trends across datasets analysed by EMinsight.

**Figure 4.**
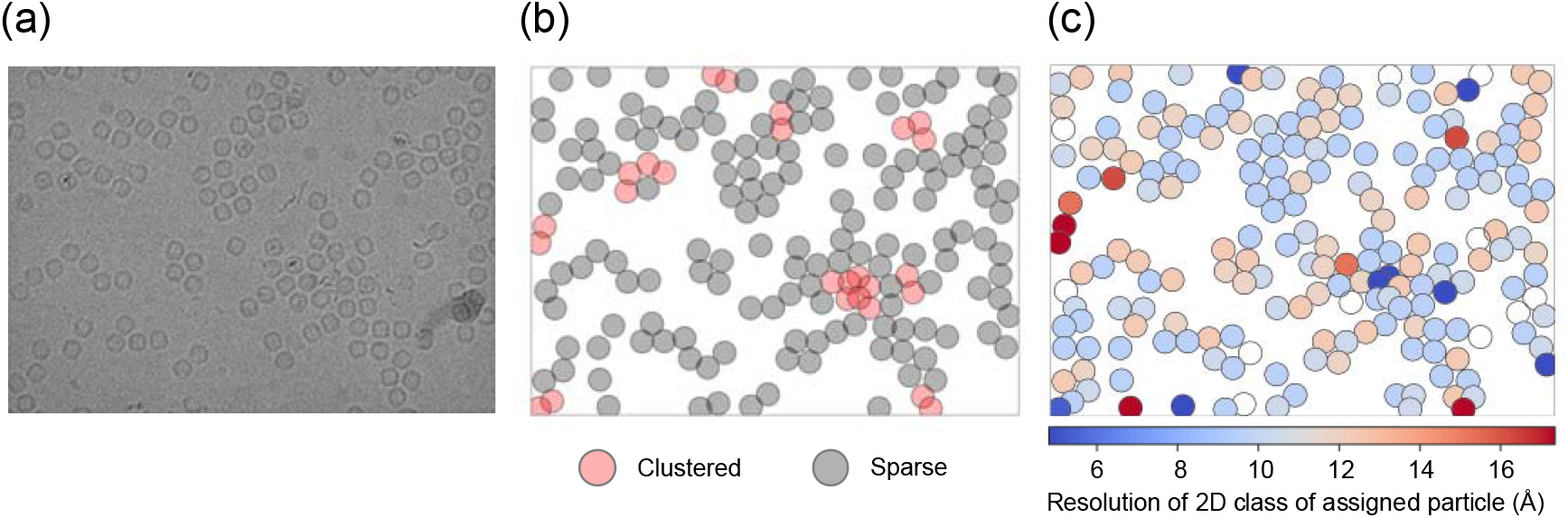
A representative set of particle coordinate analyses showing (a) micrographs, (b) the associated picks where clustered particles are represented as red and (c) the resolution of 2D classes to which particles within a micrograph are assigned.

**Figure 5.**
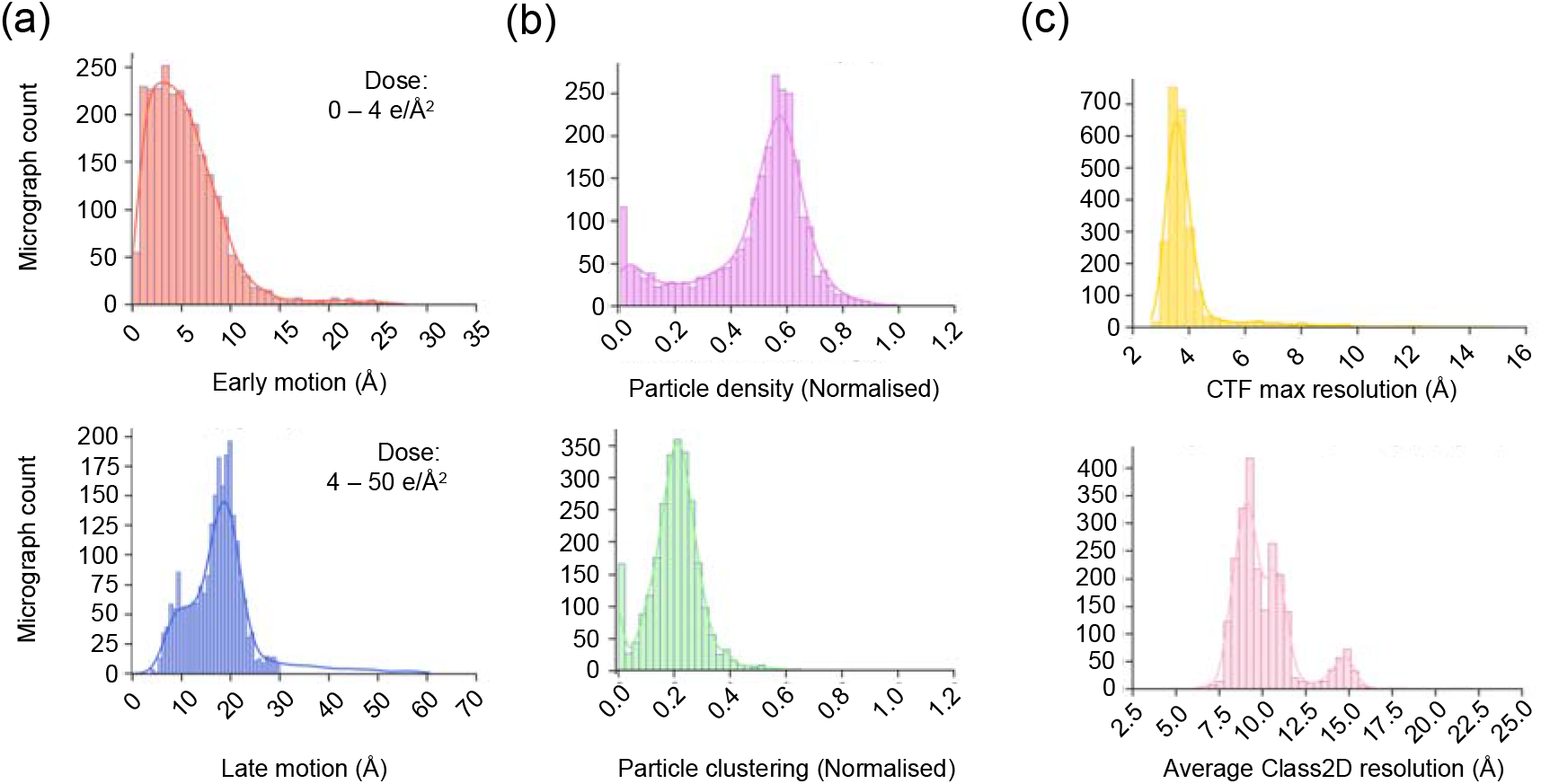
A representative set of analyses of pre-processing results prepared by EMinsight showing (a) specimen motion early (top) and late (bottom) during an image acquisition with a total dose of 50 e/Å^2^, (b) normalised particle density for a particle with a measured diameter of 136 Å (top) and degree of particle clustering where particles are closer than 109 Å (bottom), (c) micrograph CTF max resolution (top) and the mean resolution of the 2D classes to which particles from a micrograph are assigned to.

The particle quality metrics are stored along with specimen motion and CTF max resolution for each micrograph of the dataset in an attempt to concisely represent the quality of the whole SPA experiment, as shown in 0. The pre-processing results captured by EMinsight will reflect the quality of the specimen but may also be influenced by the instruments performance in recording high quality information of the specimen, and so should be considered on a case-by-case basis. Where the collated data CSV outputs of EMinsight connect pre-processing results with instrument, experimental and derived metadata describing an SPA experiment, it may be possible to investigate if particular instrument configurations are deterministic in pre-processing pipeline outcomes. EMinsight thus provides a way for the microscope operator, facilities or data scientists to rapidly quantify and identify sessions that were experimentally successful and provides a framework for investigating how the instrument and specimen may together influence experimental success.

### 3.4. Analytical outcomes linked to specimen location

Every micrograph name is stored in by EMinsight in a way that allows the identification of all low magnification images used to locate that target on the specimen. Additionally, each micrographs characteristics from the preprocessing pipelines are exposed by EMinsight and stored in the context of their locating atlas, grid square, hole and hole acquisition area image. Thus, micrograph characteristics from preprocessing pipelines can be displayed for specific areas of the grid. Metadata and preprocessing results may be analysed at various hierarchical levels of the imaging experiment. For instance, aggregating all data would reveal the overall behaviour of the specimen with respect to the entire grid or atlas. Data can then be separated at the level of the grid square or the data acquisition area within a hole of the specimen support. At the level of the grid square, the user may learn about the variability in the sample, due to large variations in vitreous ice properties from the plunge freezing process. Whereas at the level of shots per hole a user may learn about the behaviour of the specimen inside the hole of the specimen support. Analysing the behaviour of the specimen inside the hole may be particularly interesting if a user could learn from this analysis to retarget their data collection. Figure 6 shows an analysis of particle behaviour and quality from all micrographs of a dataset but separated into those exposures taken at the top or bottom of the foil hole. The resolutions of the 2D classes to which individual particles are assigned ultimately suggests the particles are of equivalent quality in both target areas, despite differences in packing density and clustering. However, for datasets exhibiting pathological problems in particle distribution in holes it is expected that this type of analysis could be beneficial to retarget the data collection to collect higher quality data.

**Figure 6.**
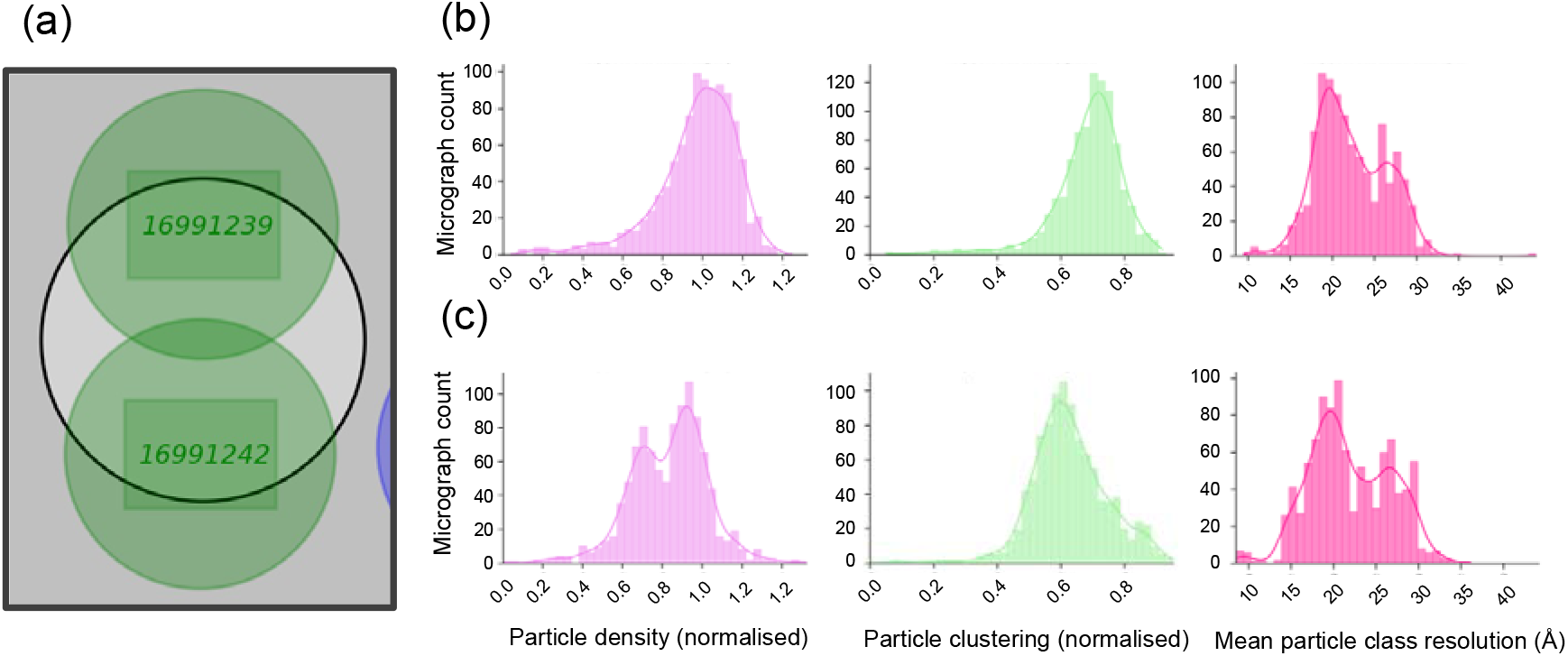
Analysing experiment performance in the context of instrument location metadata reveals trends in the particle density, aggregation and quality of particles in an SPA specimen. (a) Depicts the two exposure areas targeted for collection and for the top exposure (b) the statistics for all exposures taken in that hole location are shown, in comparison to the bottom exposure (c) the statistics for all exposures taken in that hole location.

### 3.5. Instrument setup, performance and experimental outcome described in one database

The data structure at eBIC separates experimental visits by instrument, year, user group and visit number as shown in Table 3. Due to the predictable data structure at eBIC and given the established methodology for parsing a single EPU SPA experimental metadata it is possible to systematically query every experimental visit on an instrument, for a particular user group or across the whole user programme. All these data can then be aggregated to analyse trends across multiple data collections.

**Table 3.**
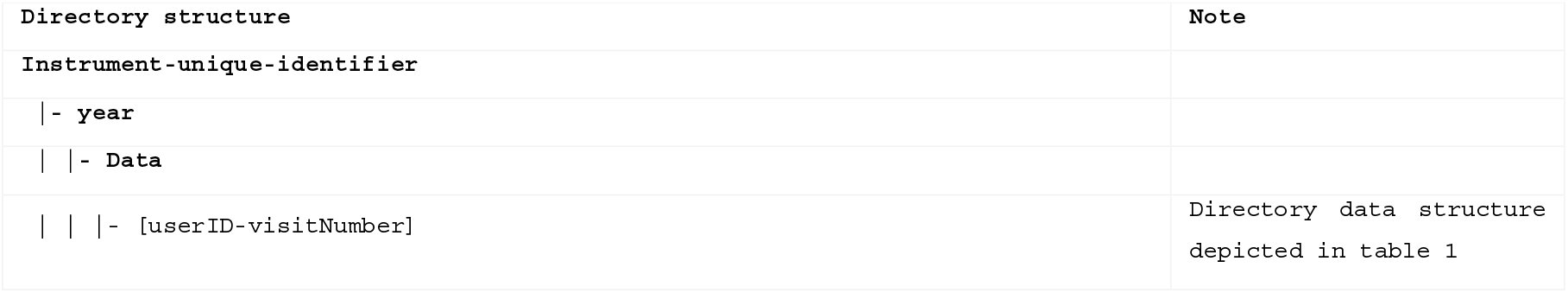
Global data structure for experimental visits at a large user facility.

Figure 7 depicts several common performance metrics recorded from SPA cryoEM experiments. EMinsight performs data reduction to produce a single value, either as an average or a max/min value to describe a collection session. Whilst within each data collection itself, these metrics will vary for each micrograph, histograms of the single reduced values aim to provide a rapid overview of the performance of an instrument or user programme. The CTF best resolutions are most commonly less than 3 Å, many sessions are subject to high levels of specimen motion but the trend is towards data exhibiting motion less than 200 Å. At the level of the specimen, most datasets are collected with micrographs exhibiting particles at less than ideal occupancy (optimal would equal a normalised value of 1), and most commonly datasets on average have low levels (∼20%) of particles found to be clustered but all datasets suffer from a degree of particle clustering.

**Figure 7.**
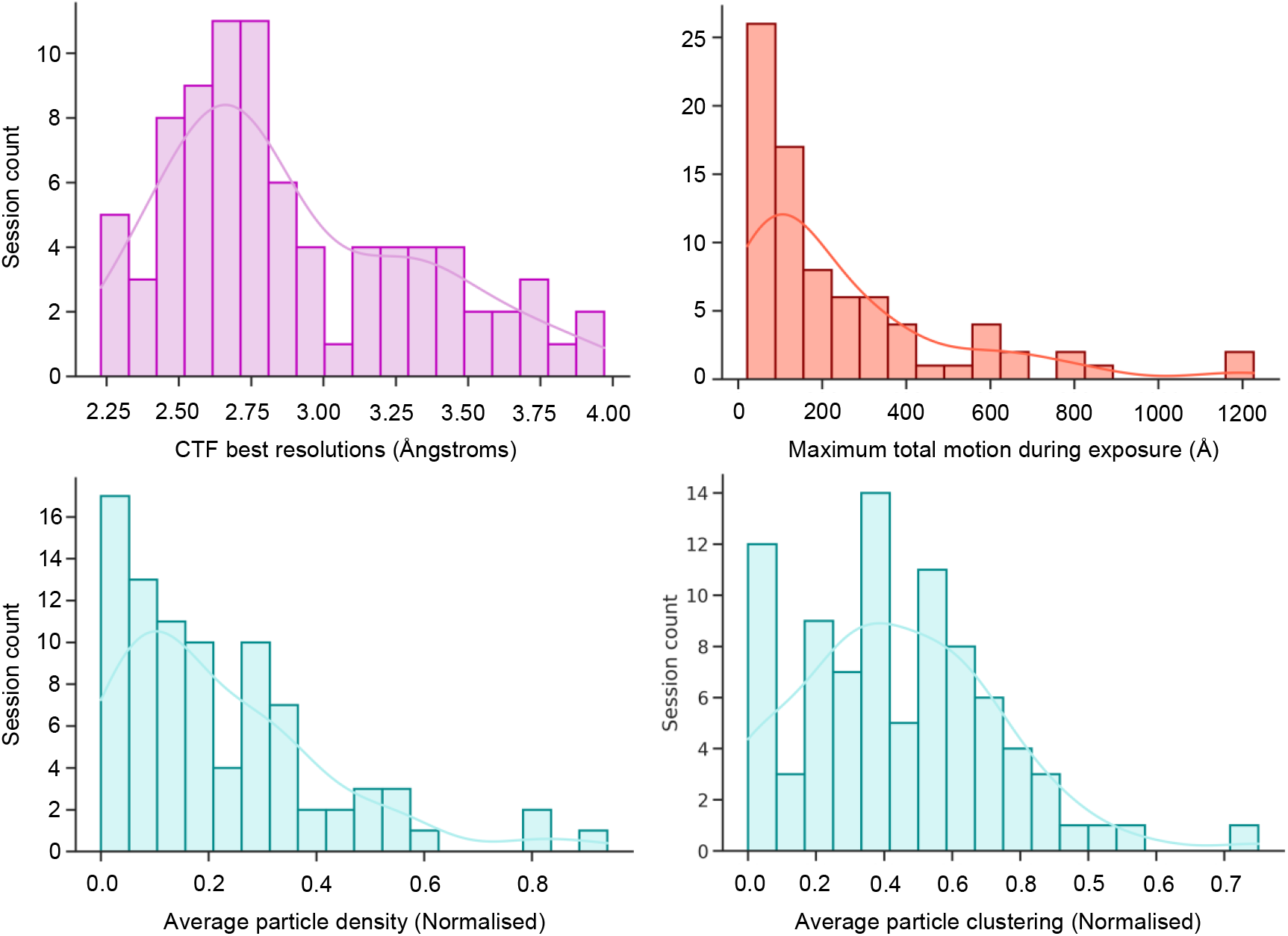
Several common performance metrics for cryoEM SPA experiments assessed across several months of a cryoEM Krios user programme.

### 3.6. Deposition ready data

EMinsight reinforces the concept for deposition configuration files, as has been suggested by other software packages (Gómez-Blanco *et al*., 2018, Kimanius *et al*., 2021). From metadata that is captured and exposed by EMinsight as reported in Table 2, a subset is extracted and stored with a view to be useful for deposition of SPA data and 3-dimensional reconstructions to the archives.

EMinsight prepares a single JSON file with much of the data fields necessary to populate the ‘Experiment: Microscopy’ sections of an EMDB archive entry. A checksum file is produced as a method to verify the Integrity of the data within the deposition JSON file. Table 4 shows the deposition file fields generated by EMinsight.

**Table 4.** An example of the fields populated by EMinsight in an example deposition file with extended metadata expected to be beneficial to complement those currently required by an EMDB archive deposition.

|  |  |
| --- | --- |
| Microscope | TITAN52334150 |
| epuversion | 3.0.0-446.0 |
| date | YYYY/MM/DD |
| eV | 300000 |
| Mag | 81000 |
| Apix | 2.3 |
| nominal_defocus_min_microns* | -1 |
| nominal_defocus_max_microns* | -3 |
| spot_size | 5 |
| C2_micron | 50 |
| Objective_micron | 100um |
| beam_diameter_micron | 1.1 |
| Collection | AFIS |
| number_of_images | 254 |
| grid_type | HoleyCarbon |
| available_squares | 62 |
| collected_squares | 3 |
| average_foils_per_square | 140 |
| hole_size_micron | 1.2 |
| hole_space_micron | 1.3 |
| shots_per_hole | 2 |
| total_dose_eA2 | 34.6 |
| fraction_dose_eA2 | 0.6 |

### 3.7. ML-ready data

As previously described, each micrograph that is assigned quality metrics is stored in a way that associates that micrograph with the lower magnification images in the hierarchical image structure taken to target that micrograph (see 2.5.1). EMinsight then introduces the concept that metadata describing high magnification images could be used as quality labels for lower magnification images that would have been collected prior to high magnification acquisition. This could be leveraged to allow the application of machine learning techniques in recognising features in low magnification images that lead to high quality data acquisition in micrographs.

Further, EMinsight’s automatic collection of metadata sufficient for the experimental section of an EMDB deposition may represent one potential avenue towards interfacing with EMDB (Lawson *et al*., 2016) and EMPIAR (Iudin *et al*., 2023), perhaps in future deposition procedures. Table 4 reports the fields that are collected by EMinsight. Some of these fields are not currently stored in archive depositions but these extended metadata fields may prove relevant for increasing the descriptive power of a EMDB deposition entry. All in all, EMinsight could represent the type of end user tool required to minimise the barrier to data submission to archives, whilst also maximising opportunities for data reuse through metadata deposition carefully considered within recommended frameworks (Sarkans *et al*., 2021). We envisage this to greatly benefit future machine learning projects relying on open-access, accurate and descriptive metadata of database entries.

## 4. Discussion

The widespread use of software-based solutions for interaction with the electron microscope has improved the efficiency of data collection and enabled almost continuous automated instrument utilisation. Despite these advances, ther’’s still a need for effective record-keeping of instrument and experimental setups, especially given the complexity of SPA experiments. A single particle analysis (SPA) cryoEM experiment is often fully described in the metadata output by the instrument itself, however these are not practically human readable. EMinsight addresses this by converting intricate metadata into human-readable reports, aiding in the documentation of SPA experiments and potentially assisting with deposition to archives.

The capturing of instrument configuration and experimental outcomes to derive calculated parameters then presents the opportunity for systematic analyses into factors affecting instrument performance. EMinsight offers insights into instrument performance, by drawing from recorded metadata but further can make inferences about the quality of the experiment and specimen by analysing pre-processing metadata. By capturing location data, analysis at various levels is enabled, such as GridSquare or exposures within a hole, to identify issues with cryoEM SPA specimens. These tool help correlate instrument configurations with experimental outcomes, expecting to be useful microscope operators, EMinsight users and facility managers in evaluating session success.

For deposition, EMinsight facilitates the recall of experiment details, which could be leveraged for populating metadata fields in structural biology archives like EMDB and EMPIAR. When TFS EPU has been used for data collection, EMinsight can produce files that could support parsing of metadata in preparation for deposition, in a concept analogous to harvesting data from structure determination applications in x-ray crystallography (Yang *et al*., 2004, Potterton *et al*., 2018). The development of automatic deposition workflows for cryoEM, possibly involving archive deposition APIs is anticipated to benefit from the deposition files produced by softwares such as EMinsight but also potentially in workflows from other softwares such as Scipion (Gómez-Blanco *et al*., 2018) or CCP-EM (Burnley *et al*.). However, the heterogeneity in data collection softwares, processing pipelines and the potential for existing local procedures in metadata capture and storage into laboratory information management systems already having been applied, increases the complexity in creating a unified system for metadata capture and automatic deposition. EMinsight is then representative of what is possible but must be considered as an example of what could be done rather than a final solution to this problem, which ultimately will require coordination from major instrument manufacturers, software developers and database developers.

Altogether, EMinsight represents a tool that gathers and relates instrument configuration, experimental outcomes, and analytical outcomes in concise reports for immediate and historical analysis of SPA data collection sessions. It could equally be adapted as a standalone tool or be incorporated into systems that feedback on cryoEM SPA experiments in real time. The coordinated recording of metadata of various kinds allows global analyses on the performance of instruments and the user programmes they run. As a tool that interprets and exposes the collections hierarchical image structure of a cryoEM SPA experiment, each micrograph is related to its low magnification images along with quality metrics and could be used as a precursor for training neural networks to recognise high quality collection areas from low and medium magnification images.

We envisage that software like EMinsight will incentivise the retention of metadata produced by cryoEM instrumentation performing SPA experiments or derived databases that describe how experiments were performed. This will improve the ability of scientists to more easily recall how experiments were performed and what their outcomes were as they develop structural biology projects aiming to determine the structures of macromolecules. These types of software could robustly inform downstream analytical processes of microscope configurations and metadata describing the experiment to automatically run image analyses. In particular, we envisage that microscope metadata could be extracted automatically at the point of map deposition or associated and retained throughout image processing to then be ready for automatically populating the archives and at the time of map deposition. More descriptive metatdata in the structural biology archives themselves could facilitate better understanding of the relationship between how an SPA experiment was performed and the quality of the resulting map, as well as enabling future ML applications on archived cryoEM data.

## Acknowledgements

Experimental data collection was performed under the supervision of David Owen, Éilís Bagginton and Vinod Vogirala by BAG user trainees on Krios 1,2 and 4 on proposal bi23047 using ApoF samples prepared by Peter Harrison and Claire Strain-Damerell. Peter Harrison is partially supported by the Membrane Protein Laboratory (Wellcome Trust grant 223727/Z/21/Z). We thank David Owen, Andrew Howe and Yuriy Chaban (eBIC, Diamond Light Source) for discussions. We thank Fanis Grollios and Reint Boer Iwema (Thermos Fisher Scientific) for discussions on EPU data structures. We thank Jason Van Rooyen (eBIC, Diamond Light Source) for support. We thank Matt Iadanza and Tom Burnley (CCP-EM) for discussions on preprocessing pipelines.

## Author contributions

KM conceived the EMinsight project and wrote the manuscript. KM wrote EMinsight with contributions from DH, JC and SR. PH prepared grids for data collection and assisted with data archiving. KM and PH tested EMinsight. All authors critically reviewed the manuscript.

## Data availability

A representative data structure on which EMinsight can be run to reproduce the analysis in this manuscript has been uploaded to EMPIAR under the accession code XXXXX.

## Source code availability

EMinsight is available in the repository: https://github.com/kylelmorris/EMinsight

## Supporting information

**Figure S1.**
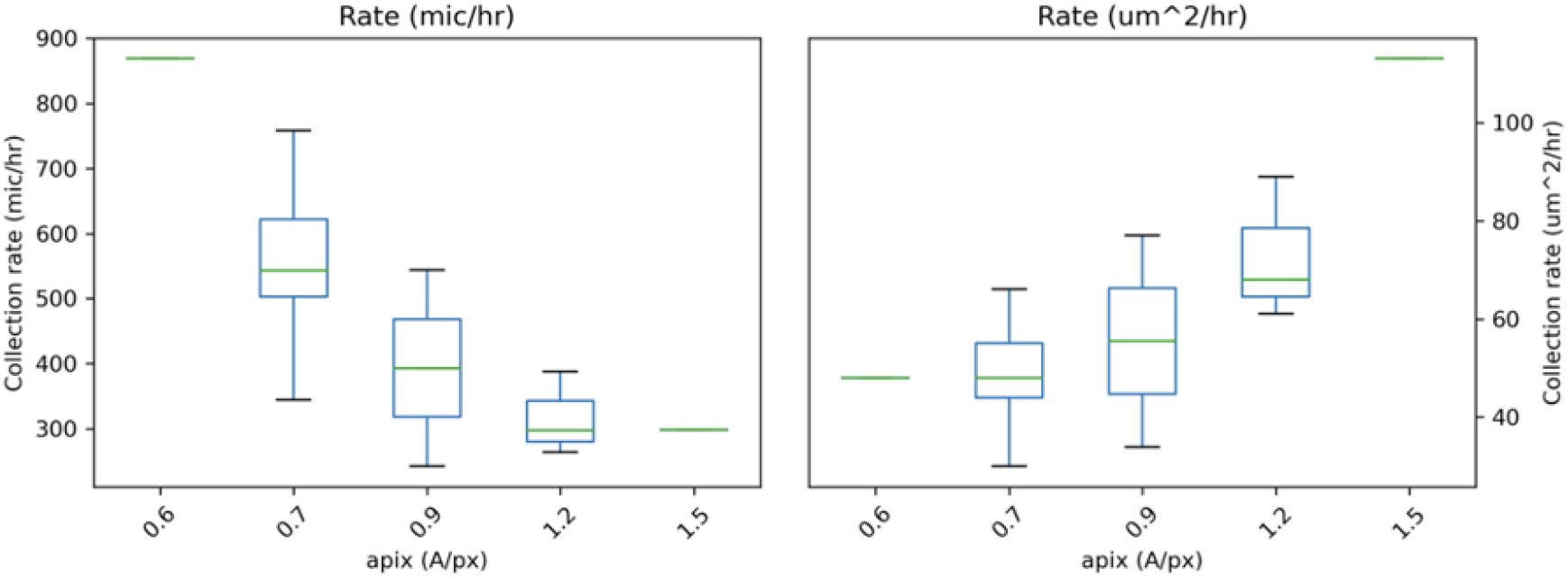
The collection performance in mic/hr and um^2^/hr for three Titan Krios microscopes with Falcon 4i and Selectris X camera/filter systems.

### S1. EMinsight session report example

The following report is produced by EMinsight describing a single SPA cryoEM experiment at the instrument performance level.

https://github.com/kylelmorris/EMinsight/blob/publication/expected_outputs/single_session/EMinsight/report/Supervisor_20230919_140141_84_bi23047-106_grid1/Supervisor_20230919_140141_84_bi23047-106_grid1_session.pdf

### S1.1. EMinsight processed report example

The following report is produced by EMinsight describing a single SPA cryoEM experiment at the analytical performance level.

https://github.com/kylelmorris/EMinsight/blob/publication/expected_outputs/single_session/EMinsight/report/Supervisor_20230919_140141_84_bi23047-106_grid1/Supervisor_20230919_140141_84_bi23047-106_grid1_processed.pdf

## S2. Supplemental files

### S2.1. EMinsight single session expected outputs

The following are representative of the files that are produced by EMinsight to store metadata on a single SPA cryoEM experiment and are referenced to produce the PDF reports

### S2.1.1. Collated data files

https://github.com/kylelmorris/EMinsight/tree/publication/expected_outputs/single_session/EMinsight/csv

### S2.1.2. Deposition files

https://github.com/kylelmorris/EMinsight/tree/publication/expected_outputs/single_session/EMinsight/dep

### S2.2. EMinsight multi session expected outputs

The following are representative of the files that are produced by EMinsight to store metadata on a multiple SPA cryoEM experiments

### S2.2.1. Multisession collated data files

https://github.com/kylelmorris/EMinsight/tree/publication/expected_outputs/global/csv

## Notes

https://github.com/kylelmorris/EMinsight

### Competing Interest Statement

The authors have declared no competing interest.

